# Vascular fungi alter the apoplastic purinergic signaling in plants by deregulating the homeostasis of extracellular ATP and its metabolite adenosine

**DOI:** 10.1101/2022.11.18.517145

**Authors:** Christopher Kesten, Valentin Leitner, Susanne Dora, James W. Sims, Julian Dindas, Cyril Zipfel, Consuelo M. De Moraes, Clara Sánchez-Rodríguez

## Abstract

Purinergic signaling activated by extracellular nucleotides and their derivative nucleosides trigger sophisticated signaling networks. The outcome of these pathways determine the capacity of the organism to survive under challenging conditions. Both extracellular ATP (eATP) and Adenosine (eAdo) act as secondary messengers in mammals, essential for immunosuppressive responses. Despite the clear role of eATP as a plant damage-associated molecular pattern, the function of its nucleoside, eAdo, and of the eAdo/eATP balance in plant stress response remain to be fully elucidated. This is particularly relevant in the context of plant/microbe interaction, where the intruder manipulates the extracellular matrix. Here, we identify Ado as a main elicitor secreted by the vascular fungus *Fusarium oxysporum*. We show that eAdo modulates the plant’s susceptibility to fungal colonization by altering eATP-mediated physiological immune responses, such as apoplastic pH and calcium homeostasis. Our work indicates that plant pathogens actively imbalance the eAdo/eATP levels as a virulence mechanism.

**One Sentence Summary:** The apoplastic Adenosine/ATP balance is a messenger for plant defense and can be manipulated by the fungal pathogen *F. oxysporum*.

## INTRODUCTION

Adenosine-5’-triphosphate (ATP) constitutes the energy currency of all living organisms and is the driving force of many cellular processes. In addition, it fulfills a broad range of tasks in signaling mechanisms once it leaves the cytosol and becomes extracellular (eATP). In plants, eATP contributes to root hair growth, gravitropism, cell death, and to response to abiotic and biotic stresses (*1–4*). ATP reaches the apoplast through transporters (*5, 6*) and secretory vesicles (*7*). Since eATP is involved in a broad selection of signaling processes, tight controlling mechanisms are required to regulate its concentration. These comprise apoplast facing apyrases and purple acid phosphatases that hydrolyze eATP to adenosine monophosphate (AMP) (*8, 9*). AMP is subsequently hydrolyzed by 5’ nucleotidases (5’NT) to extracellular adenosine (eAdo) (*8*) that is either taken up into the cytoplasm by the EQUILIBRATIVE NUCLEOSIDE TRANSPORTER 3 (ENT3) (*10*), or further processed by the extracellular protein NUCLEOSIDE HYDROLASE 3 (NSH3) (*11*). NSH3 removes the sugar moiety of eAdo and generates adenine (Ade), which is transported into the cytoplast by a purine permease transporter (PUP) (*12*).

Mechanical wounding of the plasma membrane leads to a high release of ATP to the apoplast (*3*), increasing eATP concentration up to 80 nM (*13*), which is sufficient to activate the purinoreceptor DOES NOT RESPOND TO NUCLEOTIDES 1 (P2K1/DORN1) also known as the LecRK-I.9 (lectin receptor kinase I.9) (K_d_ ~ 46 nM; (*4*)). Perception of eATP by DORN1/LecRK-I.9 induces cellular responses including increase of cytoplasmic Ca^2+^ and reactive oxygen species (ROS), activation of MAPK cascades by phosphorylation, and transcriptional reprogramming (*14*). Indeed, 60% of genes differentially regulated after application of exogenous ATP are differentially expressed during wounding processes (*4*). The role of eATP as a damage-associated molecular pattern (DAMP) is supported by the susceptibility of *dorn1* mutants to different pathogens and their lower response to a beneficial endophyte (*15, 16*).

In mammals, Ado is also recognized as a secondary messenger, being a key signal of immunosuppressive responses after being perceived by G-protein-coupled receptors (*17–19*). In plants, Arabidopsis *ent3nsh3* double mutant, affected in eAdo metabolism, showed increased susceptibility to the necrotroph ascomycete *Botrytis cinerea* by attenuating the eATP-induced ROS production and the upregulating of defense-related genes (*20, 21*). In addition, the beneficial root fungal endophyte *Serendipita indica* secretes ecto-5’- nucleotidases (E5’NT) that, like plant 5’NTs, are capable of hydrolyzing eATP and thereby shifting the equilibrium in the apoplast from eATP to eAdo (*16*). Together, these data suggest that the eATP/eAdo balance is relevant for fungal infection and that microbes might manipulate the apoplastic Ado levels in its favor, as they do with the apoplastic pH (pH_apo_) (*22–24*). However, the roles of eAdo and of the eATP/eAdo equilibrium in plant defense remain poorly understood.

*Fusarium oxysporum* (Fo) is one of the plant pathogenic fungi whose capacity to induce pH_apo_ changes is best studied (*23, 24*). As a microbe that mainly grows and advances through the apoplast, it represents an excellent model system to study plant-microbe molecular communication in this plant region. We thus made use of an elicitor mix that we obtained from the Arabidopsis pathogen Fo5176. These chemicals induced rapid and local alterations of the pH_apo_ and cellulose synthesis machinery similar to those observed by the fungal hyphae (*24*). Through fractionation followed by HPLC, we identified Ado as an abundant molecule in the Fo5176 elicitor mix. Our data indicate that Fo5176 increases the levels of eAdo in the apoplast to facilitate its growth in the host. Genetic, transcriptomic, and live-cell high-resolution microscopy approaches revealed that Ado alters ATP-induced plant defense responses.

## RESULTS

### Fo5176 secretes Ado that seems to counteract eATP-induced plant defense

We recently showed that a Fo5176 elicitor mix regulates the growth-defense balance in plants (*24*). To identify the molecules in the elicitor mix involved in this response, we performed a bioassay-guided fractionation, using a C18 solid phase cartridge with a 10% step gradient of a water:methanol solvent system followed by HPLC on a semiprep C18 column. This approach yielded a purified active component that we identified as adenosine (Ado) by standard 1D and 2D NMR (Table 1) and high-resolution mass spectrometry. Comparison of its retention time and mass data to a pure commercial standard, confirmed that Ado is a main component of elicitor mixes generated from *in vitro*-grown Fo5176 (Fig. 1A). We then asked whether the fungus secretes this potential new elicitor during plant infection. As the plant or fungal origin of the extracellular Ado (eAdo) present in the host apoplast cannot be distinguished, we tested the expression of genes required for hydrolysis and secretion of eAdo, *ENT* and *5’NT*, respectively, in the host and the intruder during their interaction. Both *FoENT* and *Fo5’NT* were significantly upregulated during root colonization (Fig. 1B and C), while the expression of the Arabidopsis’ homologs was not altered by the presence of the fungus (Fig. S1), indicating that Fo5176 might indeed secrete Ado to the apoplast while colonizing plant roots.

**Table 1:** NMR data of Ado. The chemical shifts of ^1^H and ^13^C found in Ado. Blanks are heteroatoms in the main chain. Selected COSY and HMBC correlations are included to demonstrate linkage.

| Atom | $\delta^{13}\text{C}$ | $\delta^1\text{H}$ | HMBC | COSY |
| --- | --- | --- | --- | --- |
| **1** |  |  |  |  |
| **2** | 152.4 | 1H 8.13 (s) | 4, 6 |  |
| **3** |  |  |  |  |
| **4** | 149.1 | - |  |  |
| **5** | 119.1 | - |  |  |
| **6** | 156.1 | - |  |  |
| **7** |  |  |  |  |
| **8** | 139.7 | 1H 8.34 (s) | 4, 5 |  |
| **9** |  |  |  |  |
| **1'** | 90.8 | 1H 5.95 (d, $J=3.9\text{Hz}$) | 4, 8, 2' | 2' |
| **2'** | 81.1 | 1H 4.52 (dd, $J=6.3, 3.9\text{ Hz}$) | | 1', 3' |
| **3'** | 77.1 | 1H 4.22 (dd $J=6.3, 3.8\text{ Hz}$) | | 2', 4' |
| **4'** | 87.7 | 1H 3.93 (q $J=3.8\text{ Hz}$) | | 3', 5' |
| **5'** | 62.4 | 1H 3.61 (m), 1H 3.50 (m) |  | 5', 4' |

**Figure 1:**
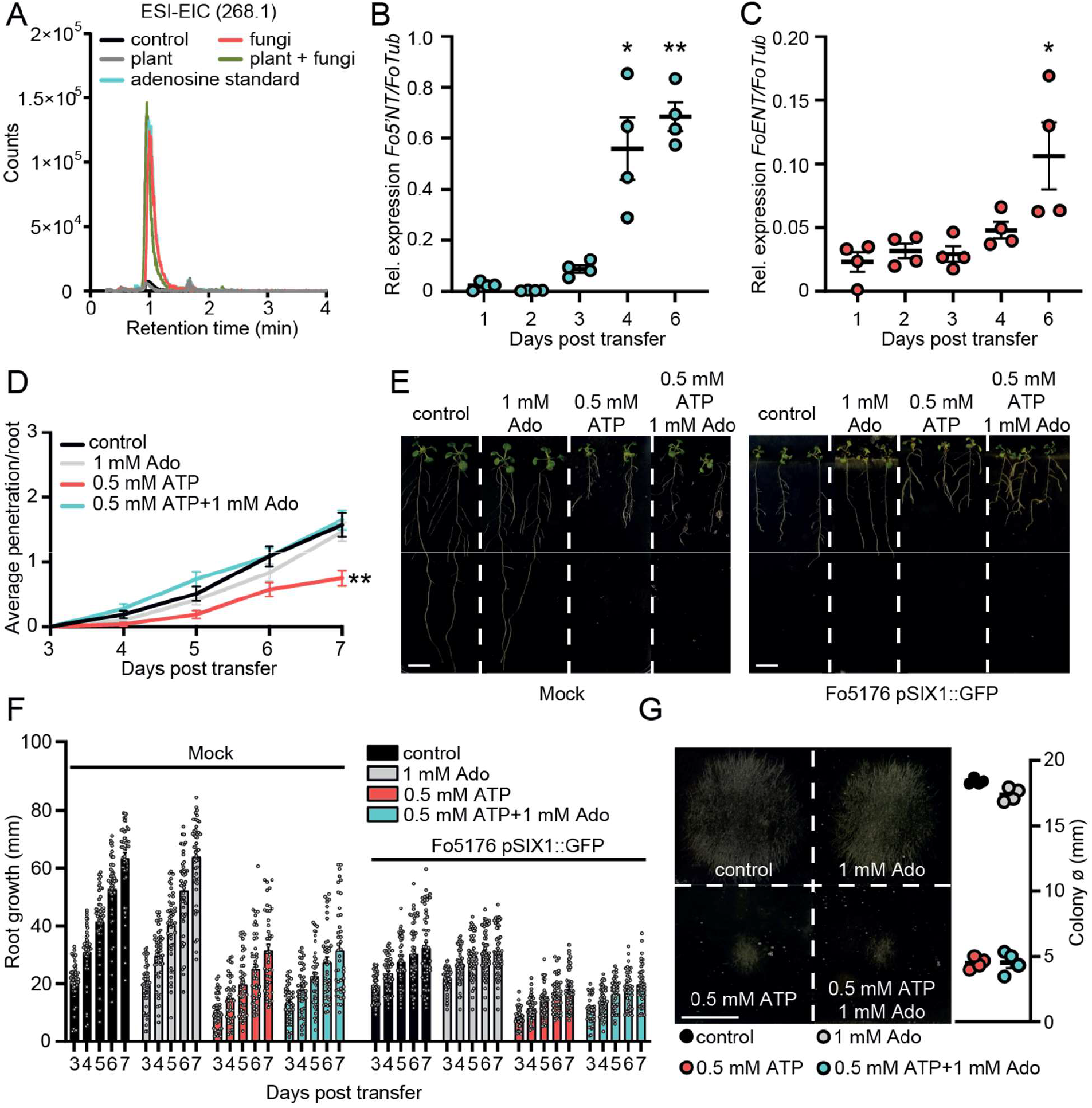
Enhanced apoplastic Ado counteracts eATP-induced reduction of fungal penetration on root vasculature. **(A)** Overlayed LC-MS extracted ion chromatograms of blank (red), adenosine (green), plant and fungus (blue). Overlayed LC-MS extracted ion chromatogram of blank (black), 250 ng adenosine (blue), fungi (red), plant (gray), plant and fungi (green) (**B)** and **(C)** *Fo5’NT* (B) and *FoENT* (C) expression relative to *FoTub* in hydroponically-grown Arabidopsis roots at various days post treatment (dpt) with Fo spores. Values are mean ± SEM, N ≥ 20, Welch’s unpaired t-test: (B) 1 dpt vs. 4 dpt: * *P*-value ≤ 0.05, 1 dpt vs. 6 dpt: ** *P*-value ≤ 0.01; (C) 1 dpt vs. 6 dpt: * *P*-value ≤ 0.05. **(D)** Cumulative Fo5176 pSIX1::GFP root vascular penetrations in wild-type (Col-0) seedlings at different days post-transfer to to plates containing ½ MS (control) and 1mM Ado and/or 0.5mM ATP. Values are mean ± SEM, N ≥ 52 from three independent experiments. RM two-way ANOVA with Tukey post-hoc test on control vs. 0.5 mM ATP: *P* ≤ 0.001 (treatment), *P* ≤ 0.001 (time), *P* ≤ 0.0001 (treatment x time). Significant differences compared to control at 7 dpt are indicated on the graph (Tukey test); statistics of remaining time points are summarized in table S2A. **(E)** Representative images of Col-0 seedlings at 7 dpt to mock (left) or Fo5176 pSIX1::GFP (right) plates. Scale bar = 1 cm. (**F)** Root growth of plants as shown in (E) at different days post transfer to mock or Fo5176 pSIX1::GFP-containing plates. Values are mean ± SEM, N ≥ 52 roots from three independent experiments, RM two-way ANOVA *P* (treatment, time, treatment x time): control vs 500 μM ATP (≤ 0.0001, ≤ 0.0001, ≤ 0.0001); control vs 500 μM ATP + 1 mM Ado (≤ 0.0001, ≤ 0.0001, ≤ 0.0001); control infected vs 500 μM ATP infected (≤ 0.0001, ≤ 0.0001, ≤ 0.001); control infected vs 500 μM ATP + 1 mM Ado-infected (≤ 0.0001, ≤ 0.0001, ≤ 0.0001). (**G)** Colony diameters of Fo5176 grown for 4 days on plates containing ½ MS (control) and 1mM Ado and/or 0.5mM ATP. Values are mean ± SEM, N = 4, Welch’s unpaired t-test control vs 0.5 mM ATP: **** *P*-value ≤ 0.0001; control vs. 0.5 mM ATP + 1 mM Ado: **** *P*-value ≤ 0.0001. Scale bar = 1 cm.

Considering the biochemical relation between Ado and ATP and the reported role of eATP in plant immunity (*2, 14*), we tested the putative influence of eAdo on ATP-induced plant defense. Thus, we first investigated if the plant response to Fo5176 is eATP-dependent by exposing the plants to different concentrations of ATP (10 μM to 500 μM) while infected by Fo5176, as described previously (*24, 25*). Indeed, 300 - 500 μM ATP significantly reduced Fo5176 vascular colonization (Fig. S2A). To assess the effect of eAdo on ATP-induced plant defense, we exposed the roots to 500 μM ATP and Ado in equimolar to doubled concentrations of ATP (Fig. S2B). Plants treated with 1 mM Ado and 500 μM eATP were indistinguishable from control plants regarding vascular penetrations by Fo5176 (Fig. 1D and S2B). Ado on its own did not have any detectable effect on fungal vascular penetration under our experimental conditions, indicating that Ado plays an important role in the plant eATP equilibrium (Fig. 1D). Importantly, eATP-induced root and fungal growth inhibition was not recovered by the addition of Ado (Fig. 1E-G). These results indicate that the plant response to Ado is ATP-dependent and implicate a mechanism in which eAdo interferes with eATP-induced plant defense responses that is not based on plant- or fungal-growth retardation.

### Plants impaired in ATP sensing or with high eAdo/eATP levels are more susceptible to Fo5176

To further test the role of Fo-secreted Ado (Fig. 1A-C) interfering with eATP during plant-pathogen interaction, we aimed at creating fungal mutants lacking *FoE5’NT* and *FoENT*. Although more than 165 potential transformants showed successful insertion of the resistance cassette into the fungal genome, none of them were knock-out mutants of the target genes, i.e. the cassette was inserted off-target. This indicates the importance of these genes for fungal viability and the very possible lethality of *FoΔE5’NT* and *FoΔENT* mutants. As manipulating the fungal eAdo levels was not successful, we addressed the influence of eAdo/eATP on plant-pathogen interactions from the plant’s side using Arabidopsis mutants altered in eAdo levels (*ent3* and *nsh3*) or eATP sensing (*dorn1)*. DORN1 impairment caused an increased fungal vascular penetration rate (Fig. 2A), underlining the role of eATP as a DAMP in Arabidopsis-Fo5176 interaction. Double mutant *ent3nsh3* plants showed an increased susceptibility to fungal colonization, while the single mutant *ent3* was not significantly affected in its response to the pathogen (Fig. 2A). Compared to Fo5176-treated WT, *dorn1* and *ent3* plants showed significantly increased primary root growth over time, while *ent3nsh3* did not differ substantially from WT, despite its higher infection numbers (Fig. 2B and C). These data indicate that the anticipated elevated apoplastic Ado levels in *enth3nsh3* (*21*) might have a main role in enhancing plant colonization by Fo5176. To test this hypothesis, we measured both soluble Ado and ATP levels in the media of hydroponically-grown plants in control and Fo5176-infected conditions, as proxy for the levels of those molecules in the apoplast. As expected, the growth media of infected *ent3nsh3* plants showed significantly elevated Ado levels in comparison to mock treatments, which were not observed in WT or *dorn1* plants (Fig. 2D). In addition, we observed significantly lower amounts of ATP in *ent3nsh3* mock-media compared to all other tested genotypes (Fig. 2E), indicating that this mutant has a higher eAdo/eATP ratio than WT in control conditions that is preserved upon Fo5176 infection due to the increase on eAdo.

**Figure 2:**
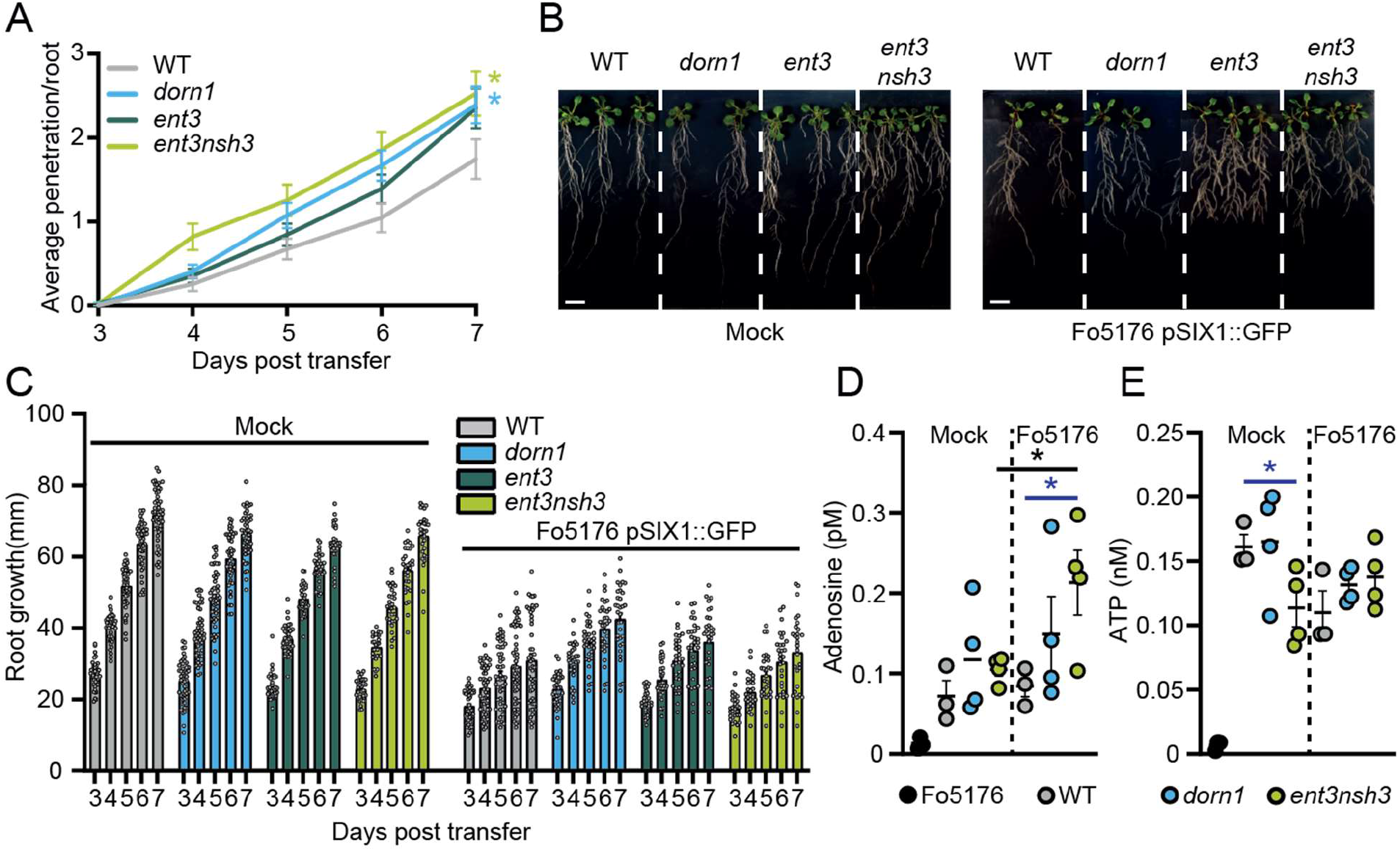
Increased extracellular Adenosine levels enhance fungal penetration rates. **(A)** Cumulative Fo5176 pSIX1::GFP root vascular penetrations in wild-type (WT; Col-0), *dorn1, ent3*, and *ent3nsh3* seedlings at different days post-transfer (dpt) to spore-containing plates. Values are mean ± SEM, N ≥ 94 from three independent experiments. RM two-way ANOVA *P* (treatment, time, treatment x time) on WT vs. *dorn1* (≤ 0.05, ≤ 0.0001, ≤ 0.05); WT vs. *ent3nsh3* (≤ 0.0075, ≤ 0.0001, ≤ 0.0061). Significant differences compared to WT plants at 7 dpt are indicated on the graph (Tukey test); statistics of remaining time points are summarized in table S2B. **(B)** Representative images of 8-day old mock or Fo5176 pSIX1::GFP infected plants as indicated in (A) at 7 days post-transfer to plates containing Fo5176 pSIX::GFP spores. Scale bar = 1 cm. **(C)** Root growth of plants indicated in (B) at different days post transfer to mock or Fo5176 pSIX1::GFP-containing plates. Values are mean ± SEM, N ≥ 79 from three independent experiments, RM two-way ANOVA *P* (genotype, time, genotype x time): WT vs. *dorn1* (≤ 0.01, ≤ 0.0001, ≤ 0.001); WT vs. *ent3* (≤ 0.0001, ≤ 0.0001, ≤ 0.0001); WT vs. *ent3nsh3* (≤ 0.0001, ≤ 0.0001, ≤ 0.05); WT infected vs. *dorn1* infected (≤ 0.0001, ≤ 0.0001, ≤ 0.0001); WT infected vs. *ent3* infected (≤ 0.05, ≤ 0.0001, ≤ 0.05). **(D)** and **(E)** Ado(D) and ATP (E) content in media from 10 days-old hydroponically-grown wild-type (WT; Col-0), *dorn1*, and *ent3nsh3* seedlings at 4 days after transfer to media with and without Fo5176 spores, and in media where Fo5176 was growing alone for 4 days. Values are mean ± SEM, N ≥ 3 biological replicates, Welch’s unpaired t-test: * *P*-value ≤ 0.05.

### eAdo increases the ATP-induced cytosolic Ca^2+^ peak

To molecularly characterize the high susceptibility of *dorn1* and *ent3nsh3* to Fo5176, we explored earlier cellular immune responses, starting with the eATP-induced cytosolic Ca^2+^ (cytoCa^2+^) peak (*4*). Employing the ratiometric cytoCa^2+^ sensor R-GECO1-mTurquoise (*26*), we first determined the minimal ATP concentration that led to a consistent increase of intracellular Ca^2+^ in the meristematic and elongation zone of Arabidopsis WT roots. 10 μM ATP were enough to consistently induce a cytoCa^2+^ peak (Fig. 3A and B), indicating that Arabidopsis roots are more sensitive to ATP than previously reported (*4*). After introgressing R-GECO1-mTurquoise into both mutant lines, we found that addition of ATP to *ent3nsh3* led to an Ca^2+^ spike 1.5 times higher than that detected in WT, indicating that an increased eAdo/eATP proportion might modulate eATP-induced stress responses (Fig. 3A-D). Consistent with the role of DORN1 as the main eATP receptor, we detected no cytoCa^2+^ peak in *dorn1* upon ATP treatment (Fig. 3E and F). Next, we investigated if the external addition of Ado can interfere with this signaling process by testing various Ado concentrations (Fig. S3A-D). While Ado did not induce any changes in the cytoCa^2+^ levels up to a concentration of 200 μM, we detected that Ado enhanced the eATP-induced cytoCa^2+^ spike when the ATP:Ado ratio was at least 1:5 (Fig. 3B). The cytoCa^2+^ spike did not further increase in response to higher eATP:eAdo ratios (Fig. S3A-D). Accordingly, the high eATP-induced cytoCa^2+^ peak observed in *ent3nsh3* did not further increase by adding Ado (Fig. 3C and D; Fig. S3E-G). These data indicate that chemical or genetic enhancement of eAdo/eATP increases the eATP-induced cytoCa^2+^ peak up to a certain eATP:eAdo concentration ratio. Moreover, we observed that eAdo could not alter the lack of response of *dorn1* to ATP (Fig. 3E and F; Fig. S3H-I), confirming that the plant response to Ado is ATP-dependent.

**Figure 3:**
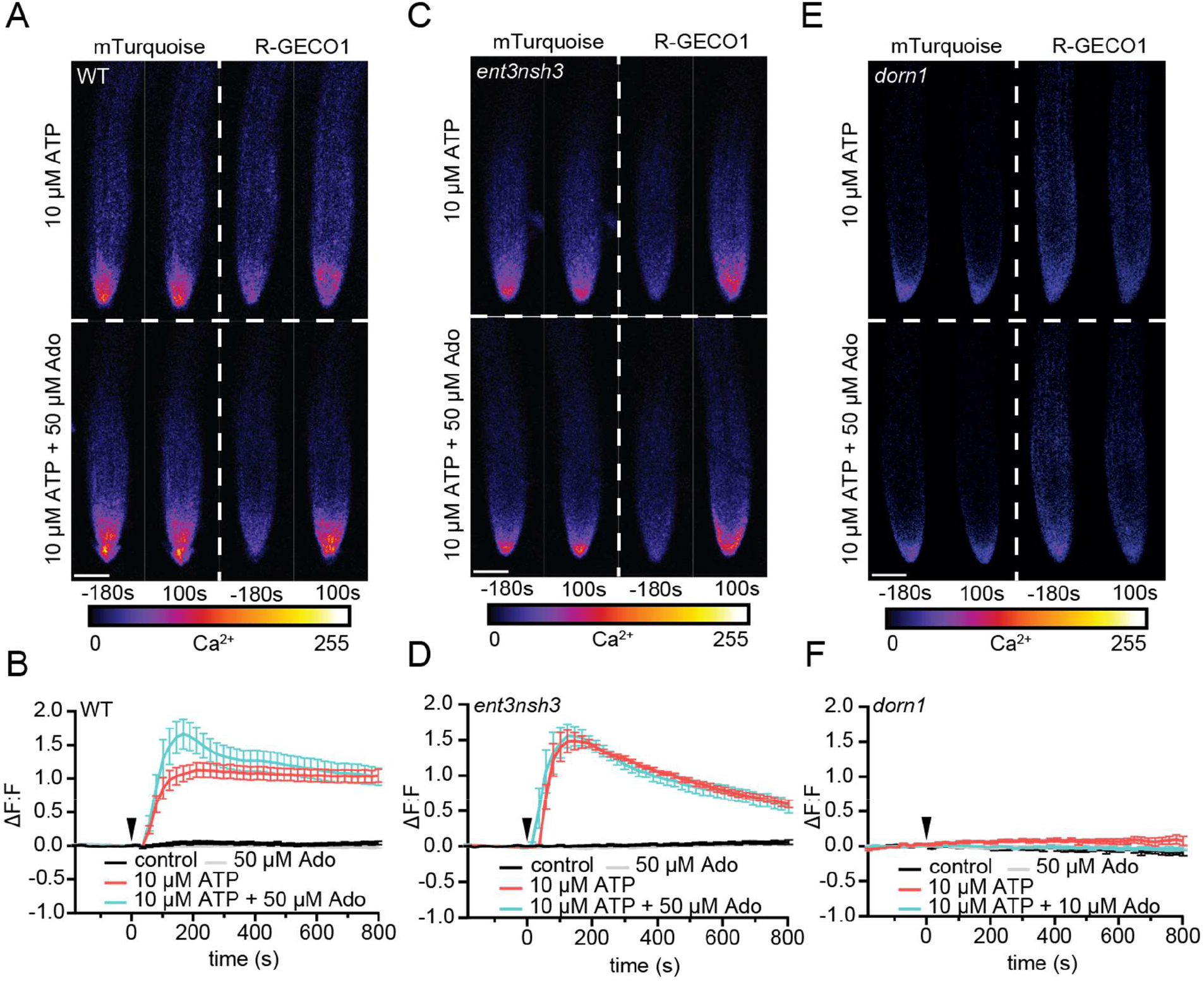
eAdo enhances eATP-induced DORN1-mediated cytosolic Ca^2+^ peak. **(A), (C)**, and **(E)** Representative images of five-days-old wild-type (WT; Col-0; A), *ent3nsh3* (C) and *dorn1* (E) roots expressing the cytoCa^2+^ sensor, R-GECO1-mTurquoise −180 s before and 100 s after being exposed to ATP (upper panels) or ATP+Ado (bottom panels). Heatmaps indicate signal intensity (arbitrary units). Scale bar = 125 μm. **(B), (D)**, and **(F)** cytoCa^2+^ in roots as in (A), (C), and (E) represented as normalized fluorescence intensity changes (ΔF:F) of R-GECO1: mTurquoise. Imaging started 180 s before either ATP or ATP+Ado was added (0 min; arrow head). Values are means ± SEM, N ≥ 18 from three independent experiments. RM two-way ANOVA *P* (treatment, time, treatment x time): **(B)** control vs. 10 μM ATP (≤ 0.0001, ≤ 0.01, ≤ 0.0001); control vs. 10 μM ATP + 50 μM Ado (≤ 0.0001, ≤ 0.01, ≤ 0.0001); 10 μM ATP vs. 10 μM ATP + 50 μM Ado (≤ 0.0001, ≤ 0.0001, ≤ 0.0001); **(D)** control vs. 10 μM ATP (≤ 0.0001, ≤ 0.0001, ≤ 0.0001); control vs. 10 μM ATP + 50 μM Ado (≤ 0.0001, ≤ 0.01, ≤ 0.0001).

### eAdo alters the ATP-induced apoplast alkalization

Exogenous application of ATP induces apoplast alkalization, as part of the fast plant response to DAMPs (*27, 28*). Hence, we investigated if, as we observed for the cytoCa^2+^ peak, eAdo also influences the ATP-dependent apoplastic pH (pH_apo_) changes. By imaging the ratiometric pH_apo_ sensor SYP122-pHusion (*24*) in WT roots, we confirmed that the apoplast alkalizes in response to the same ATP concentration required to induce a cytoCa^2+^ peak (10 μM; Fig. S4A), which we used concurrently for all further experiments. Analogous to the effect on cytoCa^2+^ levels, Ado did not affect the pH_apo_ on its own even at concentrations of 200 μM, while it altered the plant response when combined with eATP starting at 1:5 eATP:eAdo ratio (Fig. 4 and S4). eAdo concentrations up to 100 μM counteracted the eATP-induced apoplast alkalization, while 200 μM eAdo enhanced the eATP-dependent pH_apo_ peak, analogous to the cytoCa^2+^ response (Fig. 4A and B, Fig. S4A). On the other hand, *ent3nsh3* mutants showed comparable pH_apo_ response to ATP as observed in WT roots, a response that was not altered by the Ado treatment (Fig. 4C and D, S4B). These results indicate that the high eAdo/eATP ratio in *ent3nsh3* apoplast cannot alter the plant response to eATP regarding pH_apo_ changes but block the effect of exogenous Ado. In this context it has to be highlighted that *ent3nsh3* mutants already show an elevated apoplastic pH under physiological conditions (pH = 6.00) in comparison to WT (pH = 5.54) (Fig. S4D). Unexpectedly, *dorn1* responded to eATP with a slight, but significant, pH_apo_ decrease that was restored to control levels by eAdo (Fig. 4E and F; S4C). Our data indicate that ATP induces a DORN1-independent apoplastic acidification, which seems to be counteracted by eAdo. Moreover, *dorn1* roots also showed a more alkaline apoplast than WT under control conditions, as detected in *ent3nsh3* (Fig. S4D), hinting at a disturbed proton homeostasis in both mutants.

**Figure 4:**
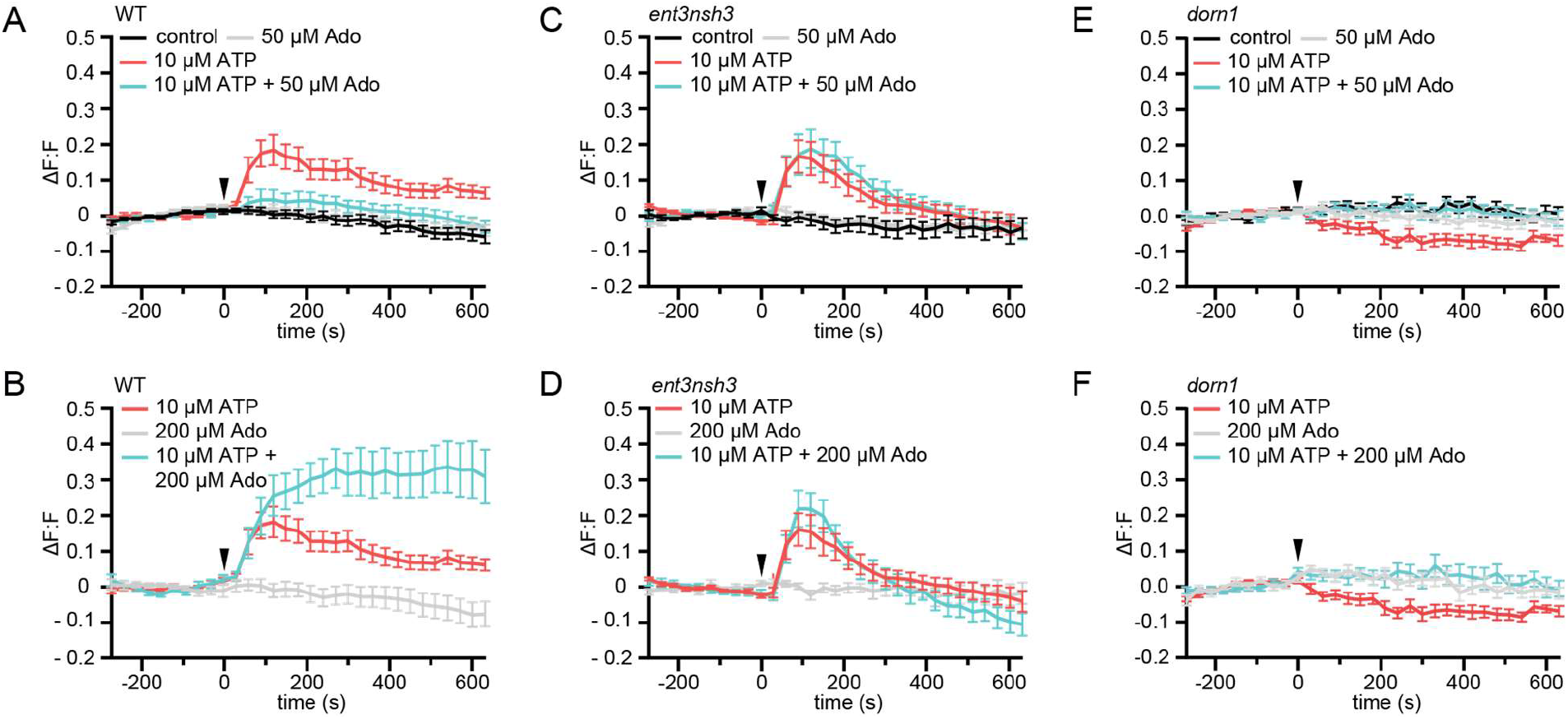
eAdo alters eATP-induced DORN1-mediated apoplast alkalization. **(A), (B), (C), (D), (E)**, and **(F)** Apoplastic pH over time in roots expressing the pH_apo_ sensor SYP122-pHusion represented as the relative signal compared to the averaged baseline recorded prior to treatments (ΔF:F).Imaging started 270 s before either ATP or ATP+Ado was added (0 s; arrow head). Values are mean ± SEM; N ≥ 12 seedlings from three independent experiments. RM two-way ANOVA, *P* (treatment, time, treatment x time) on **(A)** control vs. 10 μM ATP (≤ 0.0001, ≤ 0.0001, ≤ 0.0001); 10 μM ATP vs. 10 μM ATP + 50 μM Ado (≥ 0.05, ≤ 0.0001, ≤ 0.05) **(B)** 200 μM Ado vs. 10 μM ATP + 200 μM Ado (≤ 0.05, ≤ 0.001, ≤ 0.0001); 10 μM ATP vs. 10 μM ATP + 200 μM Ado (≤ 0.01, ≤ 0.01, ≤ 0.0001); 200 μM Ado vs. 10 μM ATP (≤ 0.05, ≤ 0.001, ≤ 0.0001); **(C)** control vs. 10 μM ATP (≤ 0.05, ≤ 0.0001, ≤ 0.0001); control vs. 10 μM ATP + 50 μM Ado (≤ 0.05, ≤ 0.0001, ≤ 0.0001); ATP vs. 10 μM ATP + 50 μM Ado (≤ 0.01, ≤ 0.0001, ≤ 0.0001) **(D)** 200 μM Ado vs. 10 μM ATP (≥ 0.05, ≤ 0.0001, ≤ 0.0001); 200 μM Ado vs. 10 μM ATP + 200 μM Ado (≥ 0.05, ≤ 0.0001, ≤ 0.0001); 10 μM ATP vs. 200 μM Ado + 10 μM ATP **(E)** control vs. 10 μM ATP (≤ 0.001, ≤ 0.0001, ≤ 0.0001; **(F)** 200 μM Ado vs. 10 μM ATP (≥ 0.05, ≤ 0.001, ≤ 0.01).

### The expression of Arabidopsis defense genes in response to Fo5176 is eATP/eAdo-dependent

To further investigate the influence of Fo5176 on the activation of eATP/eAdo-dependent plant immune responses, we measured the expression of various defense-related genes upon Fo5176 infection. In agreement with the function of eATP as DAMP, *DORN1* expression increased in WT infected-roots but was significantly downregulated in *ent3nsh3* mutant plants upon Fo5176 colonization (Fig. 5). The expression of three genes previously reported to be activated in Fo5176-infected Arabidopsis roots, *WRKY45, WRKY53*, and *At1g51890 (23, 24, 29)*, followed a similar pattern as they were all upregulated in response to Fo5176 in WT plants but not in *dorn1* or *ent3nsh3* mutants (Fig. 5). These data confirm that Fo5176 induces a eATP/eAdo-dependent plant immune response that might explain the high susceptibility of *dorn1* and *ent3nsh3* mutants to the fungus.

**Figure 5:**
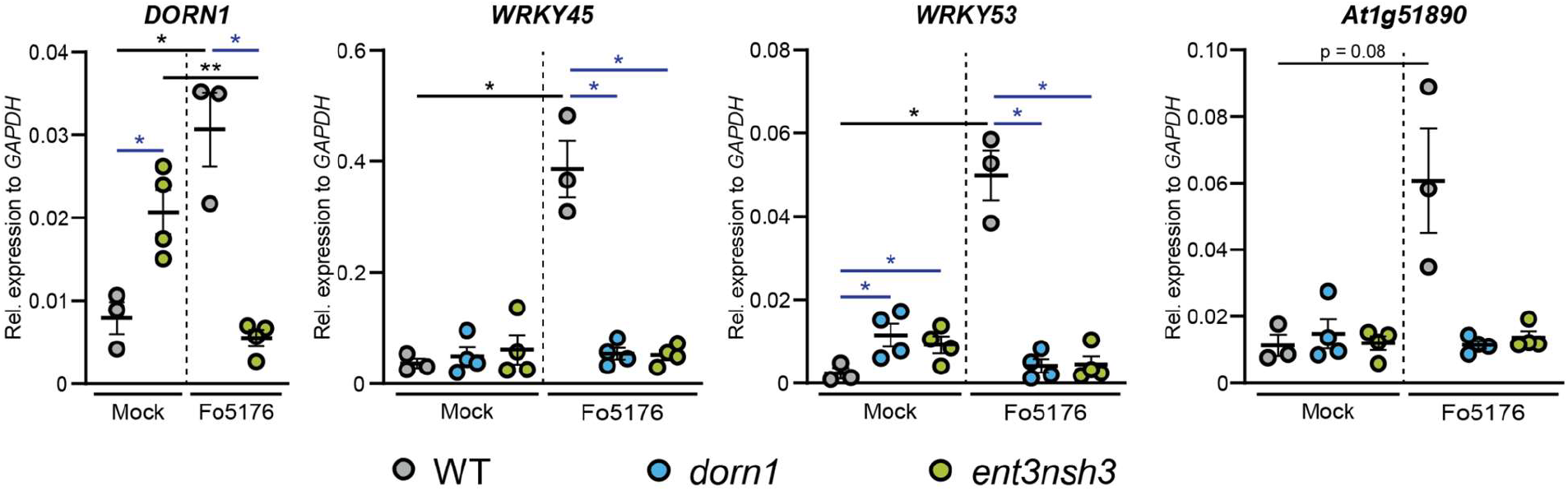
Accumulation of extracellular Ado impedes DORN1-mediated gene defense upregulation. *DONR1, WRKY45, WRKY53*, and *At1g51890* expression relative to *AtGAPDH* in WT (Col-0), *dorn1*, or *ent3nsh3* roots 4 days post-treatment with Fo5176 spores or with control media (Mock). Values are mean ± SEM, N ≥ 3 biological replicates, Welch’s unpaired t-test within each genotype in respect to their mock (black) or among genotypes (blue); * *P*-value ≤ 0.05, ** *P*-value ≤ 0.01.

## DISCUSSION

Ado is known as a key extracellular mediator of the animal immune response and its molecular activity in relation with eATP is increasingly recognized (*19, 30*). However, the knowledge of its role in plant-microbe interaction is very scarce.

In this work, we identify Ado as a main elicitor of Fo when grown *in vitro* (Fig. 1A). The transcriptional upregulation of the fungal but not of the plant eAdo producing molecular machinery during infection indicates that Fo exudes this molecule when colonizing roots (Fig. 1B and C, S1). The presence of Ado in the media during Fo colonization of Arabidopsis roots did not alter the host-microbe interaction on its own, but blocked the ATP-induced plant resistance, while not affecting fungal growth (Fig. 1D-G, S2). Hence, we deduce that eAdo interferes with the ATP-induced plant immune system activation upon fungal infection. Genetically encoded plant sensors for cytosolic Ca^2+^ levels and apoplastic pH confirmed that just 10μM ATP induces pattern-triggered immunity in Arabidopsis roots, as we could detect an immediate cytoCa^2+^ peak and apoplastic alkalization in response to this molecule (Fig. 3A-B and 4A-B). Importantly, the application of 50μM Ado further enhanced the Ca^2+^ influx response elicited by ATP, while Ado itself was inactive (Fig. 3A and B). A comparable mechanism was already discovered in oviductal ciliated cells where Ado itself is inactive but increases ATP-induced Ca^2+^ influx through the activation of the protein kinase A (*31*). The same Ado concentration efficiently blocked the ATP-induced alkalization of the root apoplast, while Ado levels above a certain threshold increased the ATP effect significantly, hinting at a possible recognition of eAdo as a MAMP by the plant immune system once a certain eATP:eAdo ratio is achieved (Fig. 4A and B). Apoplastic alkalization has been reported to be required for Fo pathogenesis (*23*) and the capacity to acidify its apoplast upon Fo contact was shown to be a positive regulator of plant defense (*24*). Therefore, our data indicate that eAdo hinders ATP-induced plant resistance by blocking ATP-induced apoplastic alkalization.

To characterize the eAdo/eATP plant signaling pathway in more detail, we studied Arabidopsis mutants altered in eATP processing and sensing, *ent3* and *nsh3*, and *dorn1*, respectively. Plants lacking the purinergic DORN1 receptor exhibited an elevated number of Fo penetrations per root compared to WT (Fig. 2A), which confirms that, as reported in other pathosystems (*14*), DORN1 is required for ATP-induced plant defense to vascular pathogens. The deficient defense responses on the transcriptional level in *dorn1* and their lack of cytoCa^2+^ changes in response to ATP +/- Ado demonstrated the role of this receptor in ATP-induced signal transduction and confirmed the inability of Ado to trigger cytoCa^2+^ peaks in the absence of ATP-signaling (Fig. 3E and F) (*4*). Accordingly, DORN1 is required for ATP-induced apoplast alkalization and its alteration when Ado is added (Fig. 4). These short term responses explain the long term effects of defective DORN1 observed in infection assays, in which Ado-altered plant defense depended on the DAMP signal eATP (Fig. 1D, 2A). On the other hand, the external application of ATP induced a slight but significant apoplastic acidification in *dorn1*, suggesting the minor involvement of other proteins in ATP-sensing (Fig. 4E and F, S4C), such as the Lectin Receptor Kinase P2K2 (*32*). Another putative candidate is the alpha subunit of the heterotrimeric G Protein, since it has been reported to be essential for ATP induced Ca^2+^ and H^+^ efflux in leaves (*28*). Both potential sensors and their role in eADo/eATP-dependent signaling should be tested in future research. The elevated pH_apo_ of *dorn1* under mock conditions might indicate a positive regulatory function of signal transducers downstream of DORN1 on H^+^-ATPases (Fig. S4D). The reason for an ATP-dependent decrease in pH_apo_ in *dorn1* mutants remains to be elucidated, although another target of eATP might be involved in that response.

Infection assays revealed an increased susceptibility of *ent3nsh3* mutants compared to WT whereas *ent3* single mutants were not significantly different from WT (Fig. 2A). We did not detect higher Ado levels in the media in contact with *ent3nsh3* roots compared to WT and *dorn1* lines (Fig. 2D), confirming previous data showing similar eAdo levels in *ent3nsh3* and WT roots grown hydroponically (*21*). Our results, though, indicate that the *ent3nsh3* mutant has a higher eAdo/eATP ratio in control media than WT plants, as it secretes comparable levels of Ado but less ATP to the media in mock conditions (Fig. 2D and E). This enhanced eAdo/eATP ratio might explain the constitutive upregulation of *DORN1* in *ent3nsh3* compared to WT (Fig. 5). *ent3nsh3* maintains its capacity to respond to Fo5176, likely because the levels of ATP and Ado/ATP in the infected-media were indistinguishable among plant lines (Fig 2D and E). Thus, we conclude that a constitutively high eAdo/eATP ratio reduces plant resistance to Fo, a hypothesis substantiated by experiments in which WT plants showed similar responses when exposed to these molecules (Fig. 1D). Furthermore, deficient defense responses on the transcriptional level in the *ent3nsh3* mutants corroborated this idea (Fig. 5). The high cytoCa^2+^ peaks detected in *ent3nsh3* in response to ATP, equivalent to what we observed in WT plants upon ATP+Ado, suggest a sufficient enrichment of eAdo/eATP in the mutant to respond to ATP (Fig. 3B and D). These high ATP-induced cytoCa^2+^ levels are not enough to activate a proper defense mechanism in *ent3nsh3*, suggesting that resistance signaling is not triggered by a high content of cytoCa^2+^ but by an increase on the levels of this cytosolic messenger (Fig. 5). On the other hand, *ent3nsh3* pH_apo_ changes in response to ATP were similar to those observed in WT, but the response was not changed by the addition of Ado, which altered the alkalization peak in control plants in a concentration dependent manner (Fig. 4A-D). This could be explained by a potential proton symporter activity of ENT3 once it starts to transport Ado into the cytoplasm (*10, 33–35*). Since Ado requires the presence of ATP to elicit pH_apo_ changes in WT plants, it is conceivable that the perception of ATP by DORN1 is required to initiate essential phosphorylation of ENT3 prior to transport, as reported for an ENT-family member in mammals (*36*). In addition, since we still detected a pH_apo_ increase in *ent3nsh3* mutants in response to ATP, we hypothesize that there is another proton-distribution-modifying component involved. Considering the alkaline apoplast detected in *ent3nsh3* under mock conditions (Fig. S4D), we anticipate that a plasma-membrane localized H^+^-ATPase might be negatively controlled by eAdo. This constitutive high pH_apo_ measured in *ent3nsh3* might explain its different response to ATP + 200 μM Ado compared to WT since the proton deficiency in the apoplast restricts the plant’s ability to further increase pH_apo_ (Fig. 4B and D; S4A and B). It also has to be taken into account that the prevalence of DORN1 in *ent3nsh3* mutants is higher compared to WT (Fig. 5), which enables an enhanced induction of downstream signals like cytosolic Ca^2+^ (Fig. 3D).

Importantly, maximum cytosolic Ca^2+^ concentrations as well as pH_apo_ peaks were detected 100 s after simultaneous application of ATP and Ado, therefore we expect Ado to prompt its effect at the plasma membrane level, like eATP (Fig. 3B and D). ATP signaling is required for plant responses to Ado, suggesting the existence of a plant Ado receptor as described in animals. Considering that AtDORN1 is not directly homologous to its mammalian counterpart, an Ado-receptor analogous to the mammalian purinergic G protein coupled receptor class is unlikely. In addition, our results indicate an immunosuppressive activity of enriched apoplastic Ado/ATP levels in the Arabidopsis-Fo5176 pathosystem. We also confirm, as reported for ΔpH (*24*), that the changes in the molecular ratio of certain molecules have a stronger effect on plant response to stress than their absolute concentrations.

## MATERIALS AND METHODS

### Plant material and growth conditions

All *Arabidopsis thaliana* lines were in Col-0 background. The pH_apo_ sensor line pub10::SYP122-pHusion, the calcium sensor line pub10::R-GECO1-mTurquoise, *dorn1-3, ent3-1* and *ent3nsh3* were published previously (*4, 21, 24, 26*). Seedlings throughout all experiments were grown upright on solid, non-buffered half MS media (pH 5.75) at 24 °C with a photoperiod of 16 h for the indicated timeframes.

### Fungal material and growth conditions

*Fusarium oxysporum* Fo5176 and *Fusarium oxysporum* Fo5176 pSIX::GFP were used throughout this study. Strain culture and storage were performed as described earlier (*37*). Fo5176 was grown in liquid half potato dextrose broth (PDB) at 27 °C for 5 days in the dark. Spores were collected by filtering the suspension through miracloth, centrifuging the filtrate at 3500 rcf, discarding the supernatant and resuspending the spores in dsH_2_O.

### Fungal elicitor mix preparation, and fractionation, and molecule identification

Fungal elicitor mix was prepared as published previously (*24, 38*) and separated via enrichment using Discovery DSC-C18 (2 g) columns (Merck) with a H_2_O/MeOH gradient from 100 % to 0 % H_2_O in 10 % steps. Fractions were bioassayed and active fractions purified to individual components via an Agilent 1100 HPLC using Zorbax SB-C18 (9.4 x 150mm) semi-prep column in a linear gradient of H_2_O/MeOH and flow rate of 5 mL/min. Individual peaks were assayed for activity. The pure active compound was characterized by standard 1D and 2D NMR experiments performed at the NMR Service of the Laboratory of Organic Chemistry at ETH Zürich. All experiments were performed using d6-DMSO in a 600 MHz Bruker NMR equipped with a 5mm probe. Data was analyzed using MestreNova 8.1 software (Mestrelab Research, Spain). LC-MS data was obtained on an Agilent 6400 LC-qTOF in scanning positive mode to produce a single signal with an m/z of 268.1044 (C10H13N5O4 calc. 268.1046 1.72 ppm) and identical retention time to an external standard of adenosine.

### Fungal transformation

PCR and complementary primers (Table S1.2) were used to generate two DNA fragments with overlapping ends (*39*). A resistance cassette containing the neomycin phosphotransferase (npTII) cloned between the *A. nidulans* gdpA promoter and the trpC terminator (*40*) was used to generate two DNA fragments promoting the homologous recombination in fungal protoplasts. Protoplasts were produced as described previously (*41*) and their transformation done as reported by (*42*).

### In-vitro growth assay of Fo5176

Freshly harvested Fo5176 spores were diluted to 10^4^ spores/mL and 15 μL of it distributed on solid half MS plates containing 1 mM Ado, 0.5 mM ATP, or both, or none (control). After four days under the plant growth conditions described above **(“**Plant material and growth conditions”), the colony diameters were measured using FIJI (*43*).

### Plant plate infection assays

Plate infection assays were performed as described earlier (*24, 25*). Ado and ATP treatment plates were generated by mixing hand-warm half MS media, 0.9 % agar, with the specific amount of stock solution. Root growth was measured using FIJI (*43*).

### Hydroponic infection assay

Hydroponic infection assays were performed as previously described (*44*). 30 seeds were grown on a foam floating on 50 mL liquid ½ MS media, pH 5.75, 1 % sucrose. After seven days the media was replaced by ½ MS without sucrose. Samples supposed to be infected were inoculated with 5*10^6^ Fo5176 spores. After the indicated days post transfer to spore-containing media, roots and fungal hyphae were harvested for subsequent expression analysis and the media was filtered. For ATP quantification media was flash frozen in liquid nitrogen, for Ado quantification it was freeze dried.

### Media ATP quantification

ATP levels in media from hydroponic infection experiments of hydroponically-grown plants were analyzed using the ATP Colorimetric/Fluorometric Assay Kit (Sigma, USA) and an Infinite M1000 plate reader (Tecan, Switzerland). Assays were done as described in the manual and ATP was detected fluorescently. All samples and standards were measured in duplicates.

### Media Ado quantification

Freeze-dried media samples from hydroponic infection experiments were resuspended in 4 mL MilliQ water. The resulting mixture was loaded onto a 100 mg Discovery DSC-18 column (Supelco, USA). The column was eluted with 1 mL MilliQ water, 1 mL 70 % MilliQ water with MeOH and finally 100 % MeOH. The resulting aqueous elution was analyzed in positive mode using an Agilent 1200 Infinity II UPLC separation system coupled to an Agilent 6550 iFunnel qTOF mass spectrometer (Agilent, USA). Compounds were separated by infecting 5 μL of sample onto a Zorbax Eclipse Plus C8 RRHD UPLC column (2.1×100 mm, 1.8 μm) held at 50°C and eluting with a linear water:acetonitrile (both modified with 0.1 % formic acid) gradient, 99 % water to 99 % acetonitrile. Mass spectral data was acquired in positive mode with an electrospray ionization source and scanning a mass range of 100-2000 m/z. Quantification was done by integrating the m/z values corresponding to Ado in MassHunter Quantitative Analysis Software and compared to a standard curve generated at the time of sample measurements.

### In-vitro growth assay of Fo5176

Freshly harvested spores were diluted to 10^4^ spores/mL and 15 μL of it distributed on solid half MS plates containing 1 mM Ado, 0.5 mM ATP or both. After four days under plant growth conditions the colony diameters were measured using FIJI.

### Gene expression analysis by real-time quantitative PCR

Freeze-dried fungal and plant material from plate infection assays respectively hydroponics was ground to powder using glass beads and a TissueLyser II (Quiagen, Netherlands). Total RNA was extracted using GENEzol™ reagent (Geneaid, Taiwan) following the manufacturer’s protocol. 1 μg of RNA was used to generate first strand cDNA using the Maxima™ First Strand cDNA Synthesis-Kit (Thermo Scientific, USA) following the manufacturer’s instructions. To amplify corresponding cDNA sequences primers (Table S1.1) (*4, 23, 24, 45–47*) were used along with Fast SYBR Green Master Mix (Thermo Scientific, USA) under following cycle conditions: 95°C for 3 min, 40 cycles of 94°C for 10 s, 58°C for 15 s and 72°C for 10 s. Two technical replicates were performed for each reaction and the reference genes *AtGAPDH* and *Foβtub* were amplified on each plate for normalization. Relative expression was analyzed using the 2^-ΔCt^ method (*48*).

### Ratiometric pH_apo_ sensor imaging

Experiments were carried out as described earlier (*24*). A Leica TCS SP8-AOBS (Leica Microsystems, Germany) confocal laser scanning microscope equipped with a Leica 10× 0.3NA HC PL Fluotar Ph1 objective or a Leica Stellaris 8 equipped with a Leica HC PL APO CS2 10x/0.40 DRY were used. pHusion was excited and detected simultaneously (Excitation: GFP 488 nm, mRFP 561 nm; Detection: GFP between 500 and 545 nm; mRFP between 600 and 640 nm). Five-day old A. thaliana seedlings expressing the sensor SYP122-pHusion grown on ½ MS + 1 % sucrose were transferred to imaging chambers as described previously (*49*) but placed on top of 1 % agarose cushions. Subsequently, the chamber was filled with ½ MS, pH 5.75. Images were collected as XYt series for 15 min with a time frame of 30 s. Image settings were kept identical throughout the experiments for each reporter line. After a recovery time of 15 minutes the experiment was started by acquiring ten images of the seedling’ roots without treatment to create a baseline of averaged relative signal. Roots were imaged from the tip including their elongation zone. The different treatments were applied in a volume of 100 μL after 300 s. ΔF:F values were calculated according to following formula: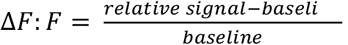. Maximal amplitudes of ΔF:F signals were obtained by averaging the maximal ΔF:F signals of all curves. To collect standard curves for the pHapo ratiometric sensor a set of nine buffers from pH 4.8 to pH 8.0 were used. Each buffer was based on 50 mM ammonium acetate. Buffer pH 4.8 comprised additionally 22 mM citric acid, 27 mM trisodium citrate. pH was adjusted with 0.01 M HCl. Buffers pH 5.2 to pH 6.4 contained 50 mM 2-(N-morpholino) ethane sulfonic acid (MES), buffers pH 6.8 to pH 8.0 were composed of 50 mM 4-(2-hydro-xyethyl)-1-piperazineethanesulfonic acid (HEPES). 1 M Bis-Tris propane was used to adjust the pH values of buffers pH 5.2 to pH 8.0. Six to eight seedlings per buffer were incubated for 15 min and imaged after transfer to microscope slides.

### Ratiometric cytoCa^2+^ sensor imaging

Imaging was done as described for the pH_apo_ sensor (*24*) with slight modifications. Five-day old *A. thaliana* seedlings expressing the reporter R-GECO1-mTurquoise (*26*) were grown on ½ MS, pH 5.75, 1 % sucrose. mTurquoise was excited with 405 nm and detected between 460 to 520 nm, R-GECO1 was excited with 561 nm and detected between 580 and 640 nm. Imaging time frame was set to 20 s. Corrective flat field images for 405 nm were acquired by using 7-Diethylamino-4-methylcoumarin (Sigma D87759-5G, 50 mg/mL in DMSO). Relative signal was calculated by dividing mean gray values of the R-GECO1 channel by the mean gray values of the mTurquoise channel. ΔF:F values and Maximal amplitude of ΔF/F signals were calculated as described for the pH_apo_ sensor.

### Statistical analyses

All statistical analyses were performed using Prism 9. Statistical methods and the resulting *P*-values are defined in the corresponding figure legends. Outlier tests were performed on datasets with. If the automatically detected fluorescent ratios of the genetic pH or Ca^2+^sensors were measured to be outside of the standard curve range, they were excluded from the analysis. Such cases could always be allocated to severe drift of analyzed roots in the analysis chamber.

## ACKNOWLEDGMENTS

Live cell imaging was performed with equipment maintained by the Scientific Center for Optical and Electron Microscopy (ScopeM, ETH Zurich) and by the Center for Advanced Bioimaging (CAB) Denmark.

## Author contributions

Conceptualization: CK and CSR. Methodology: CK, JWS, JD, CZ, CMDM, and CSR. Investigation: CK, VL, SD, JD, JWS. Funding acquisition: CK, CZ, CMDM, and CSR. Supervision: CK, CZ, CMDM, and CSR. Writing – original draft: CK, VL, and CSR. Writing – review & editing: CK, VL, SD, JWS, JD, CZ, CMDM, and CSR.

## Data and materials availability

All data are available in the main text or the supplementary materials.

## SUPPLEMENTARY MATERIALS

**Fig. S1:**
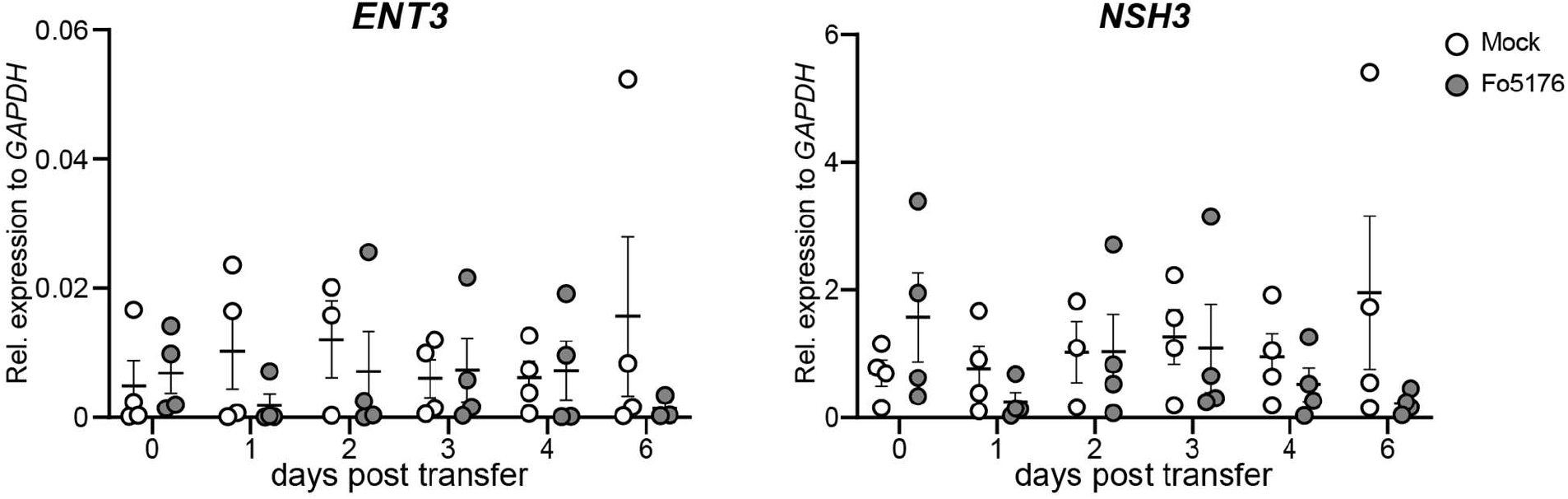
Plant *ENT3* and *NSH3* expression do not change in response to Fo5176. *ENT3* and *NSH3* expression relative to *AtGAPDH* in hydroponically-grown Arabidopsis roots at the respective days post treatment with Fo5176 spores or with control media (Mock). Values are mean ± SEM, N ≥ 3.

**Figure S2:**
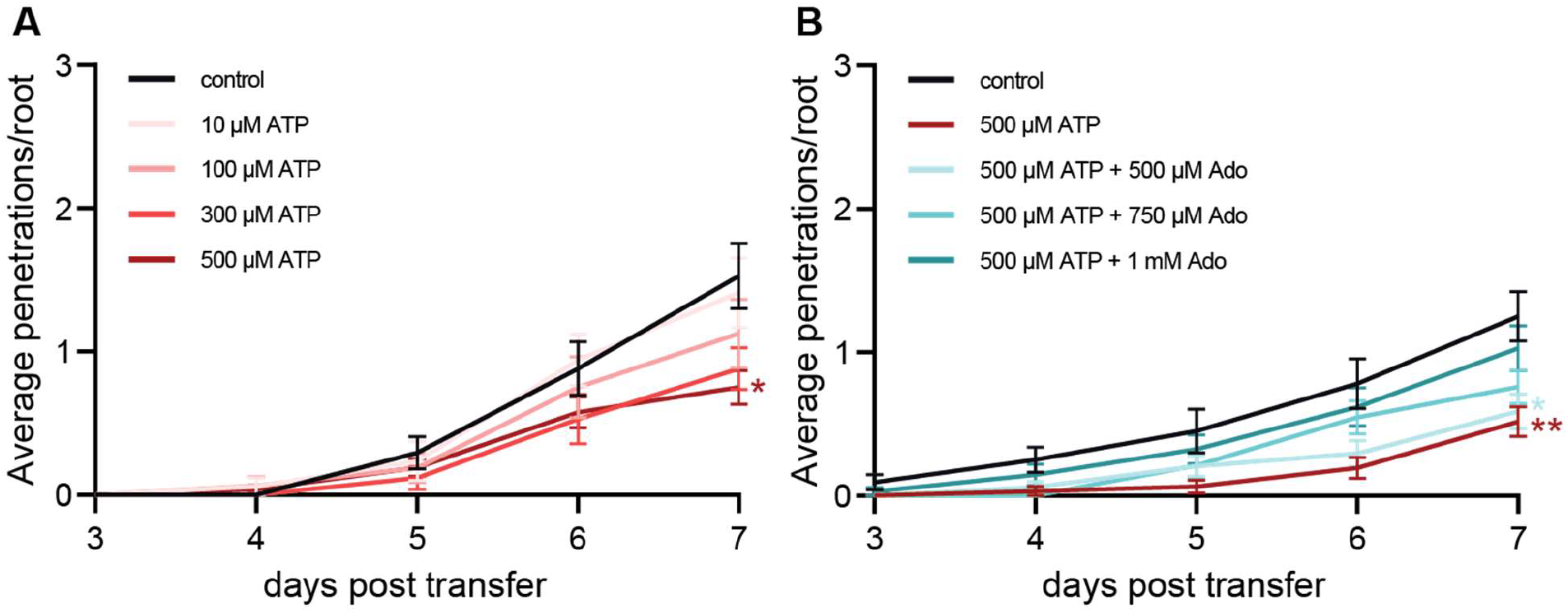
ATP-induced plant defense is counteracted by doubled concentration of eAdo. Cumulative Fo5176 pSIX1::GFP vascular penetrations per root in 8-day-old Col-0 plants at different days post transfer to plates with fungal spores alone (control), or with ATP **(A)** or ATP+Ado **(B)**. Values are mean ± SEM, N ≥ 63 from three independent experiments. RM two-way ANOVA *P* (genotype, time, genotype x time) on **(A)** control vs. 300 μM ATP (≤ 0.05, ≤ 0.0001, ≤ 0.05); control vs. 500 μM ATP (≤ 0.05, ≤ 0.0001, ≤ 0.0001); **(B)** control vs. 500 μM ATP (≤ 0.01, ≤ 0.0001, ≤ 0.0001); control vs. 500 μM ATP + 500 μM Ado (≤ 0.01, ≤ 0.0001, ≤ 0.001). Significant differences compared to control at 7 dpt are indicated on the graph (Tukey test); statistics of remaining time points are summarized in table S2C.

**Figure S3:**
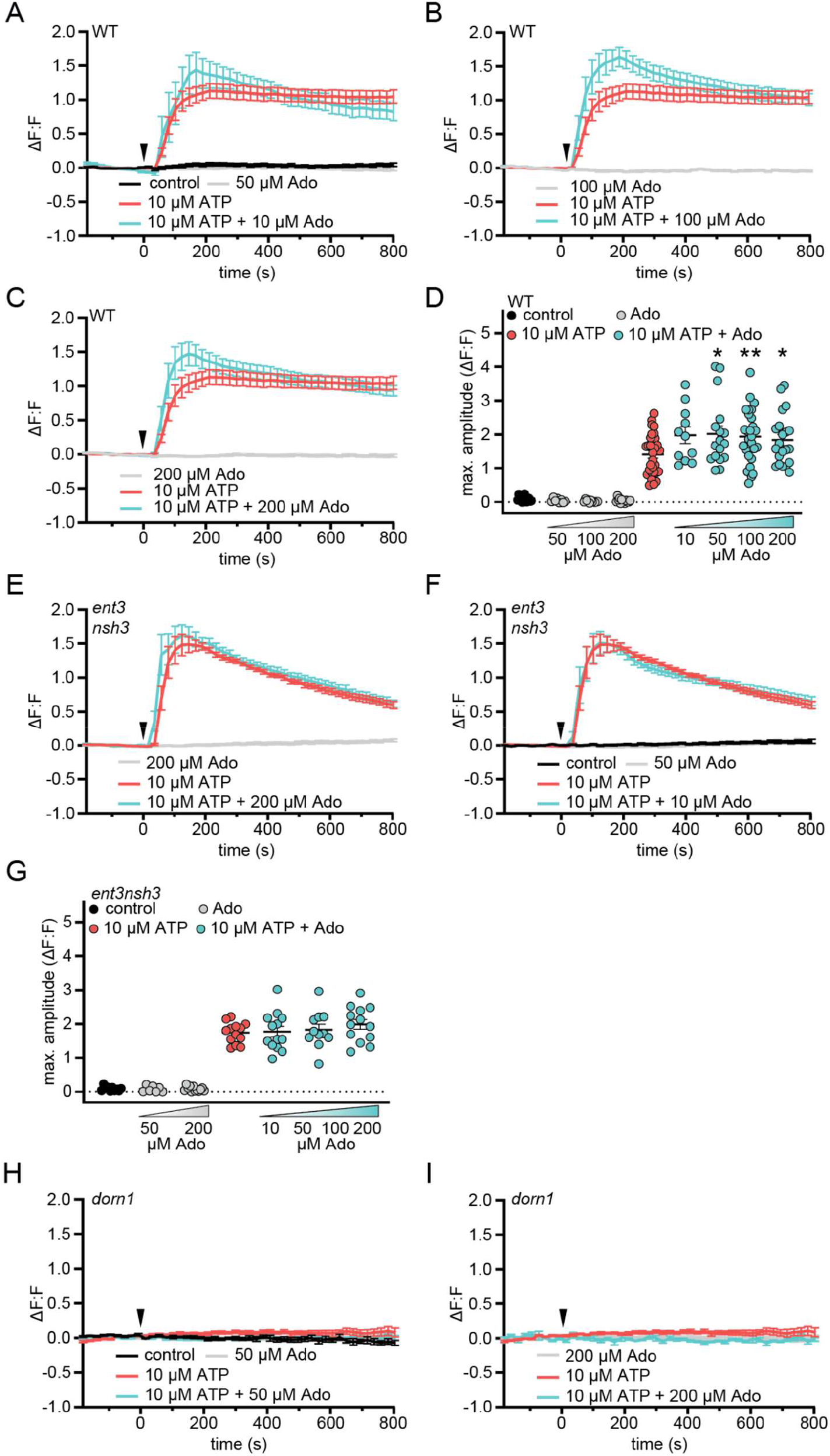
Extracellular adenosine increases extracellular ATP induced DORN1 mediated Ca^2+^ influx. Five-day-old RGECO-mTurquoise fluorometric calcium reporter line seedlings were imaged in ½ MS (−180 to 0 s)s. At 0 min, either ATP or ATP+Ado at the indicated concentrations were added ΔF:F represents the relative signal compared to the averaged baseline recorded prior to application (arrow). RM two-way ANOVA *P* (treatment, time, treatment x time): **(A)** control vs. 10 μM ATP (≤ 0.0001, ≤ 0.01, ≤ 0.0001); control vs. 10 μM ATP + 10 μM Ado (≤ 0.0001, ≤ 0.01, ≤ 0.0001); **(B)** 100 μM Ado vs. 10 μM ATP (0.0001, ≤ 0.05, ≤ 0.0001); 100 μM Ado vs. 10 μM ATP + 100 μM Ado (≤ 0.0001, ≤ 0.05, ≤ 0.0001; 10 μM ATP vs. 10 μM ATP + 100 μM Ado (≥ 0.05, ≤ 0.0001, ≤ 0.0001); **(C)** 200 μM Ado vs. 10 μM ATP (≤ 0.0001, ≤ 0.01, ≤ 0.0001); 200 μM Ado vs. 10 μM ATP + 200 μM Ado (≤ 0.0001, ≤ 0.01, ≤ 0.0001); 10 μM ATP vs. 10 μM ATP + 200 μM Ado (≥ 0.05, ≤ 0.0001, ≤ 0.0001); **(E)** 200 μM Ado vs. 10 μM ATP (≤ 0.0001, ≤ 0.01, ≤ 0.0001); 200 μM Ado vs. 10 μM ATP + 200 μM Ado (≤ 0.0001, ≤ 0.001, ≤ 0.0001; **(F)** control vs. 10 μM ATP (≤ 0.0001, ≤ 0.001, ≤ 0.0001); control vs. 10 μM ATP + 10 μM Ado (≤ 0.0001, ≤ 0.01, ≤ 0.0001); Maximum amplitudes **(D), (G)** correspond to (A) - (C) and Fig. 3 (B) (WT) respectively (E), (F) and Fig. 3 (D) (*ent3nsh3*). N ≥ 13 from three independent experiments. Welch’s unpaired t-test: * *P*-value ≤ 0.05, ** *P*-value ≤ 0.01.

**Figure S4:**
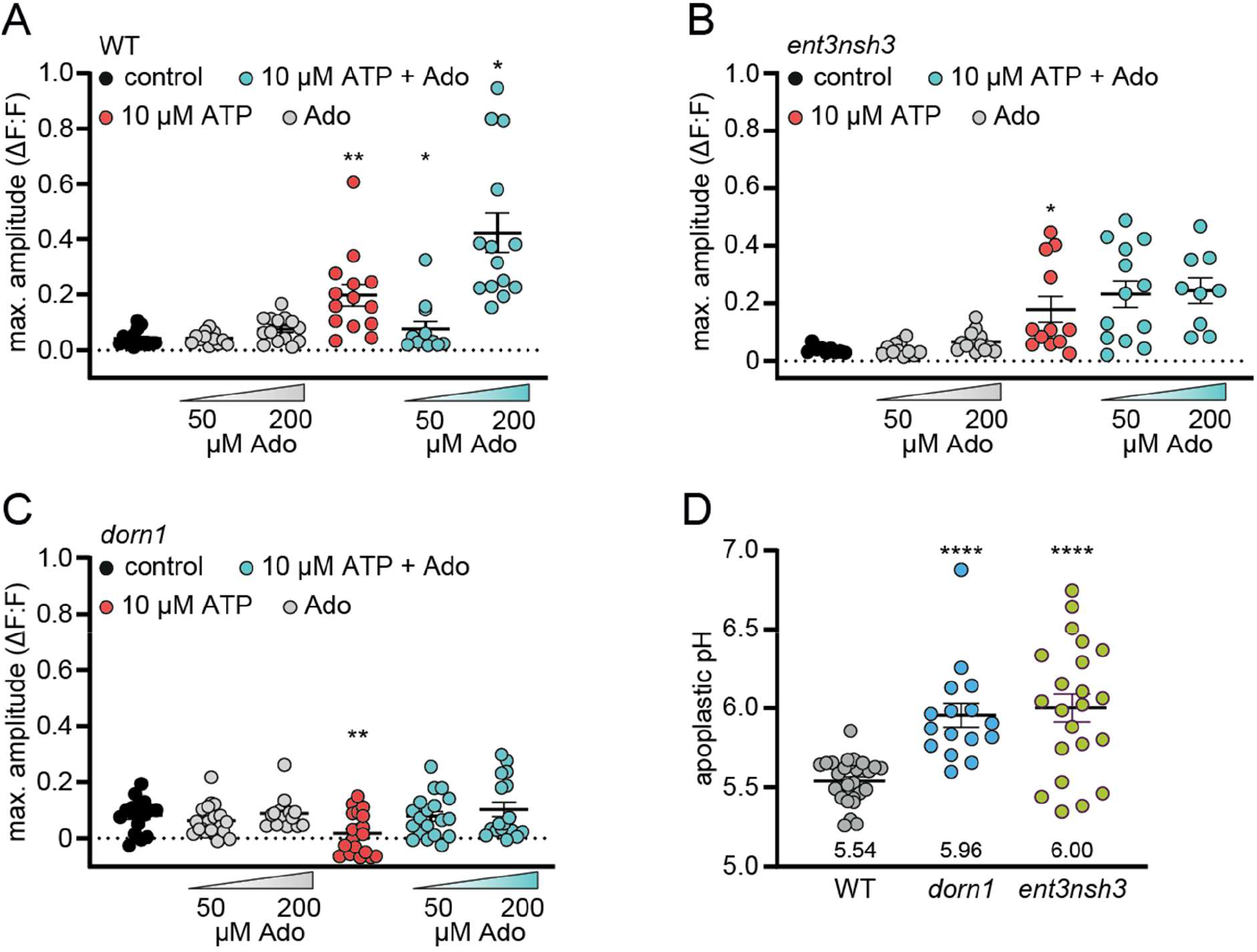
Extracellular adenosine accumulation and absent extracellular ATP receptor DORN1 elevate apoplastic pH. Five-day-old fluorometric SYP122-pHusion pH reporter lines were imaged in ½ MS pH 5.75 (−270 to 0 s) with SP8 microscope. At 0 sec, either ATP or ATP+Ado at the indicated concentrations were added. **(A), (B)** and **(C)** correspond to Fig. 4 (B), (C), (E), (F), (H), (I) and display the averaged maximal amplitude of each curve. Asterisks indicate significant differences to control (10 μM ATP) respectively to 10 μM ATP (10 μM ATP + Ado) **(D)** apoplastic pH was determined using standard curves. Values are mean ± SEM, N ≥ 16 from three independent experiments, Welch’s unpaired t-test: * *P*-value ≤ 0.05, ** *P*-value ≤ 0.01 **** *P*-value ≤ 0.0001.

**Table S1.1.**
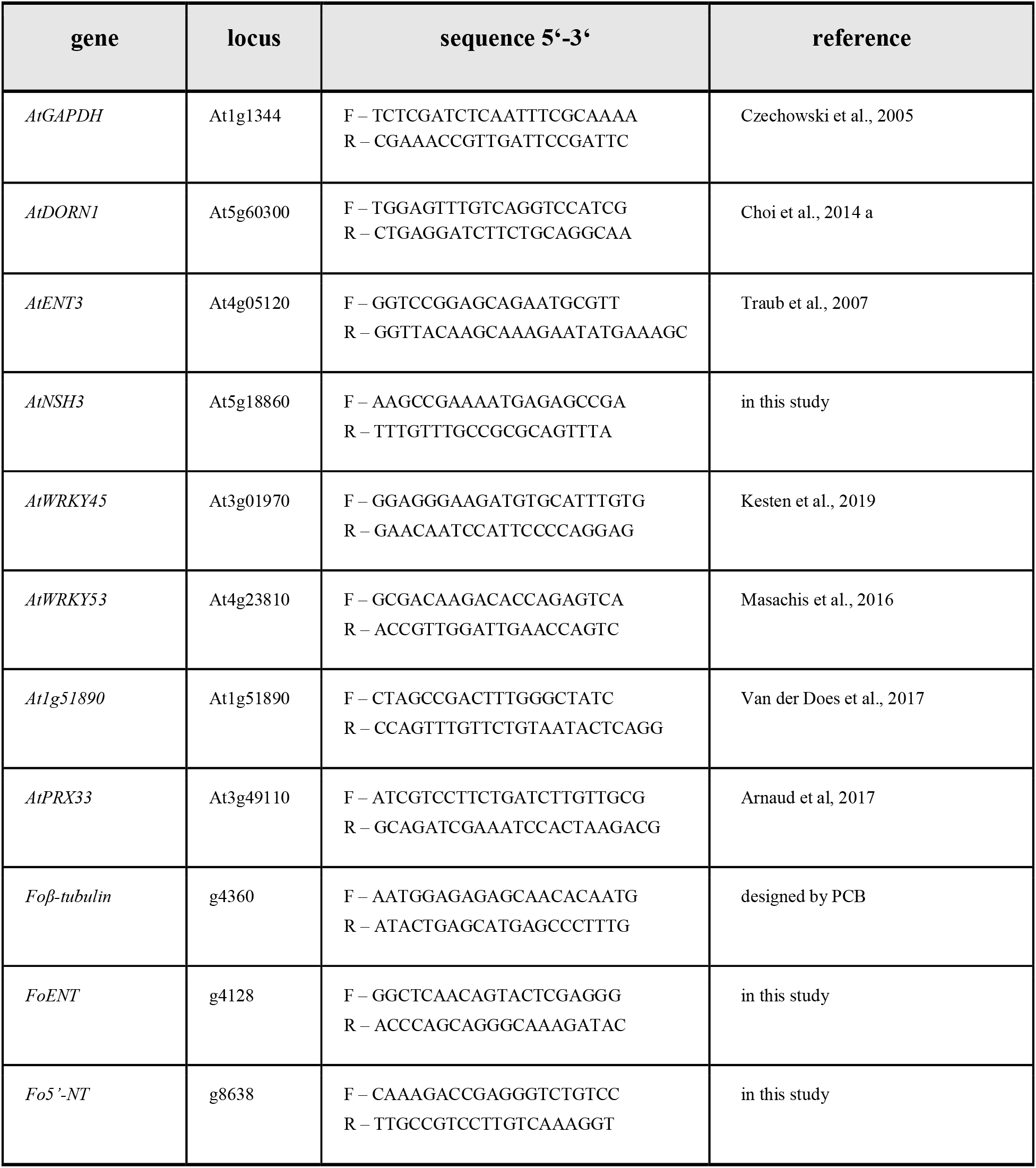
Primers used in this study for qRT-PCR.

**Table S1.2.**
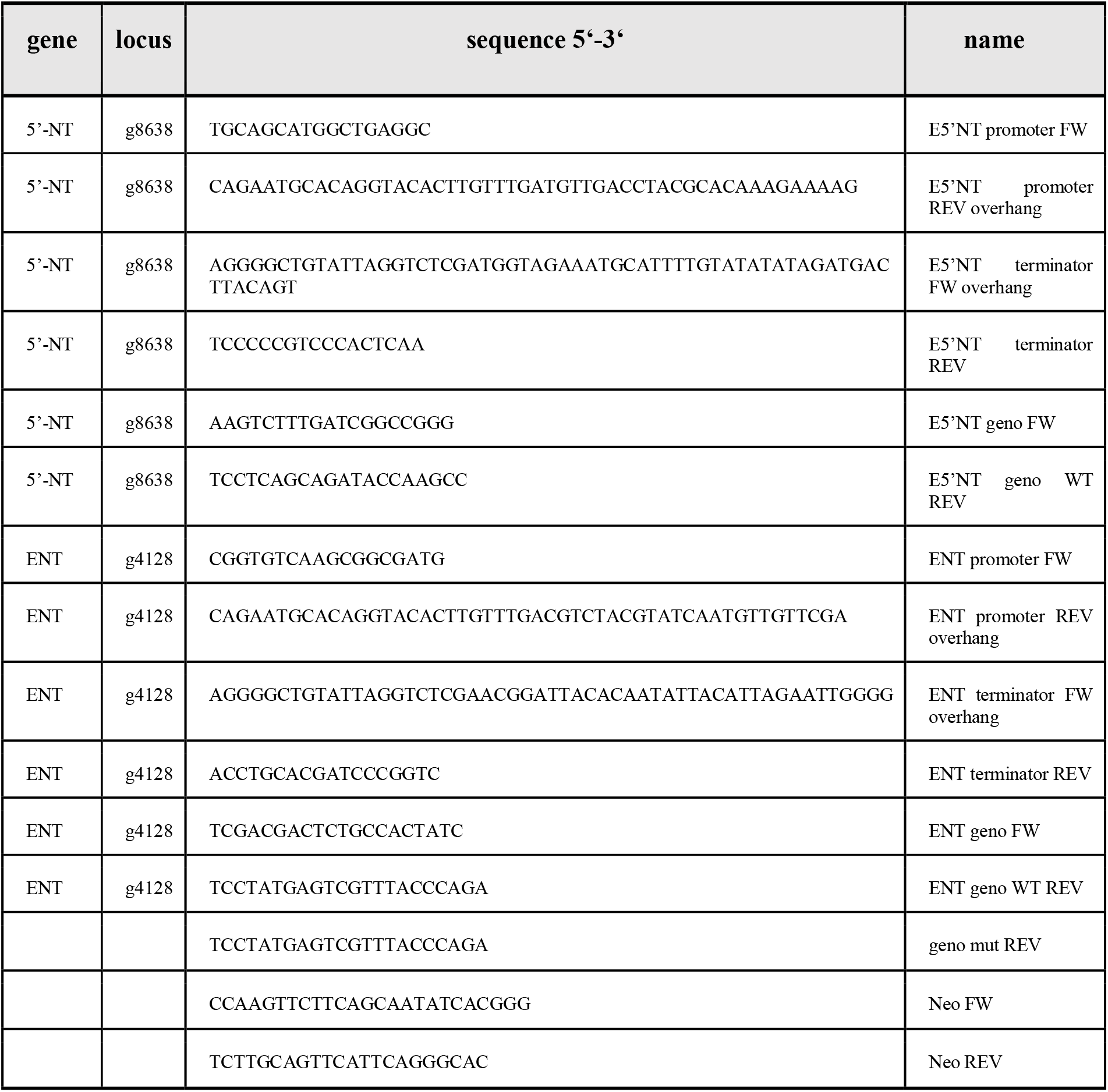
Primers used to mutagenize Fo5176.

**Tables S2.**
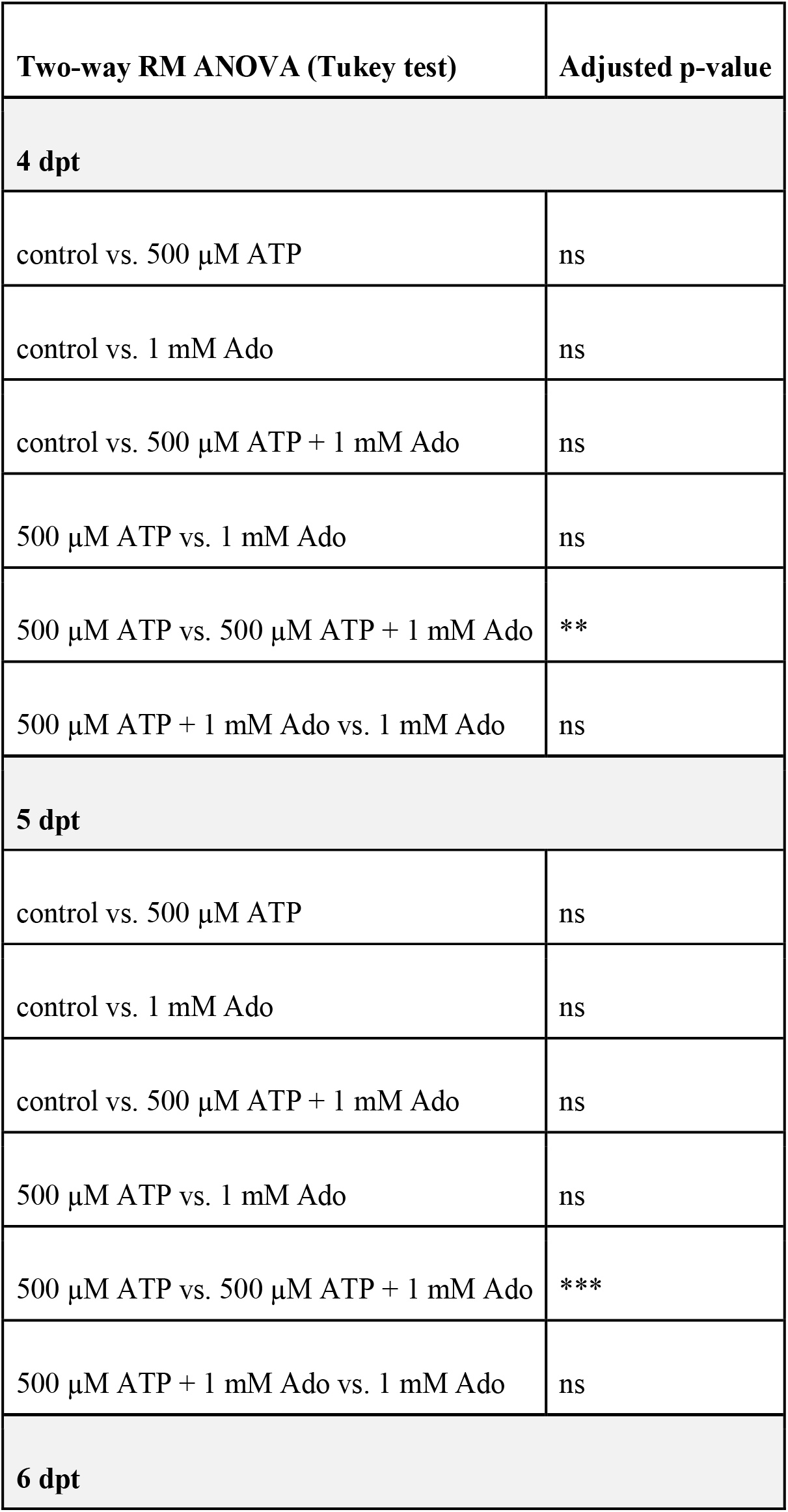
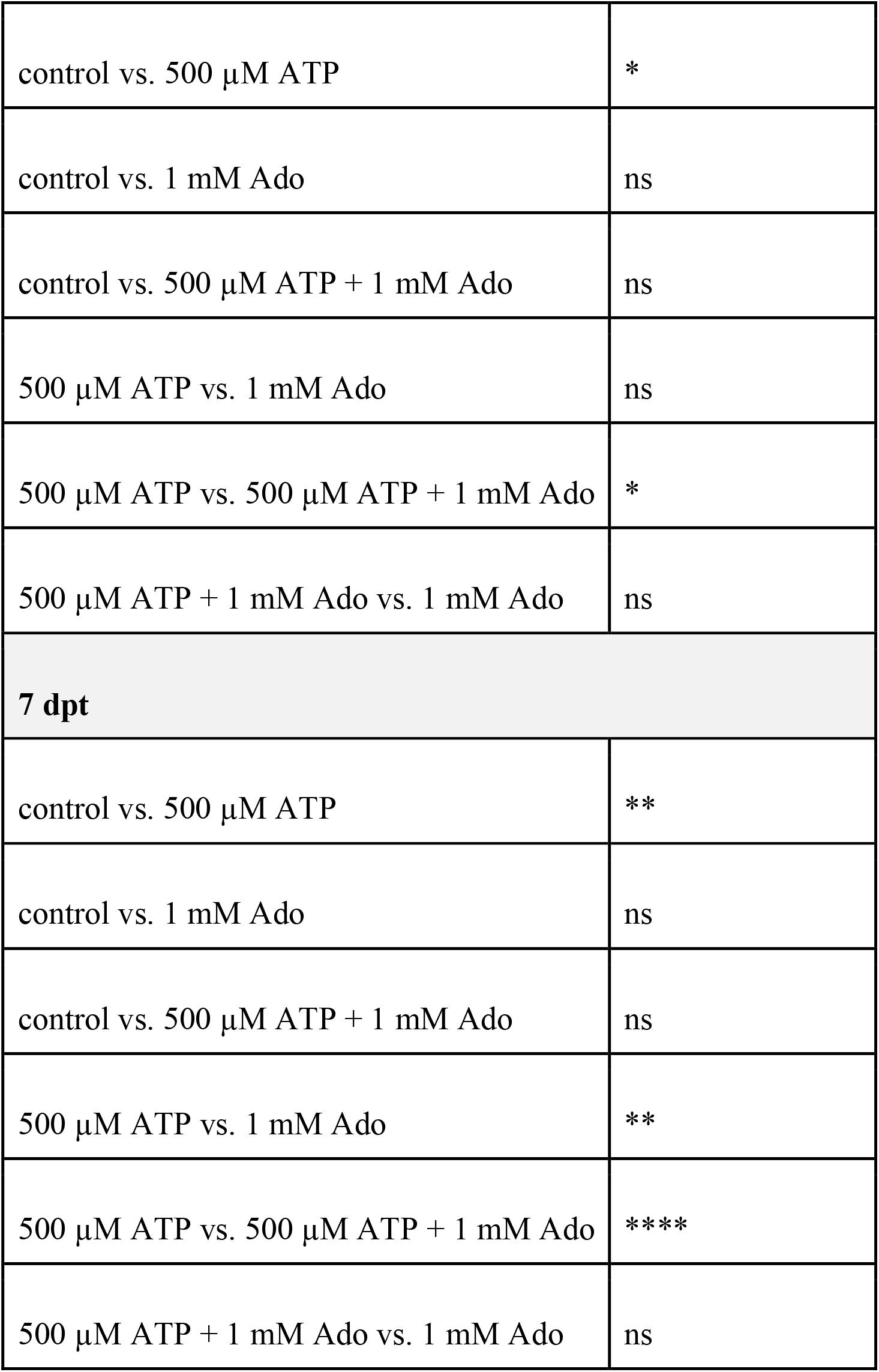
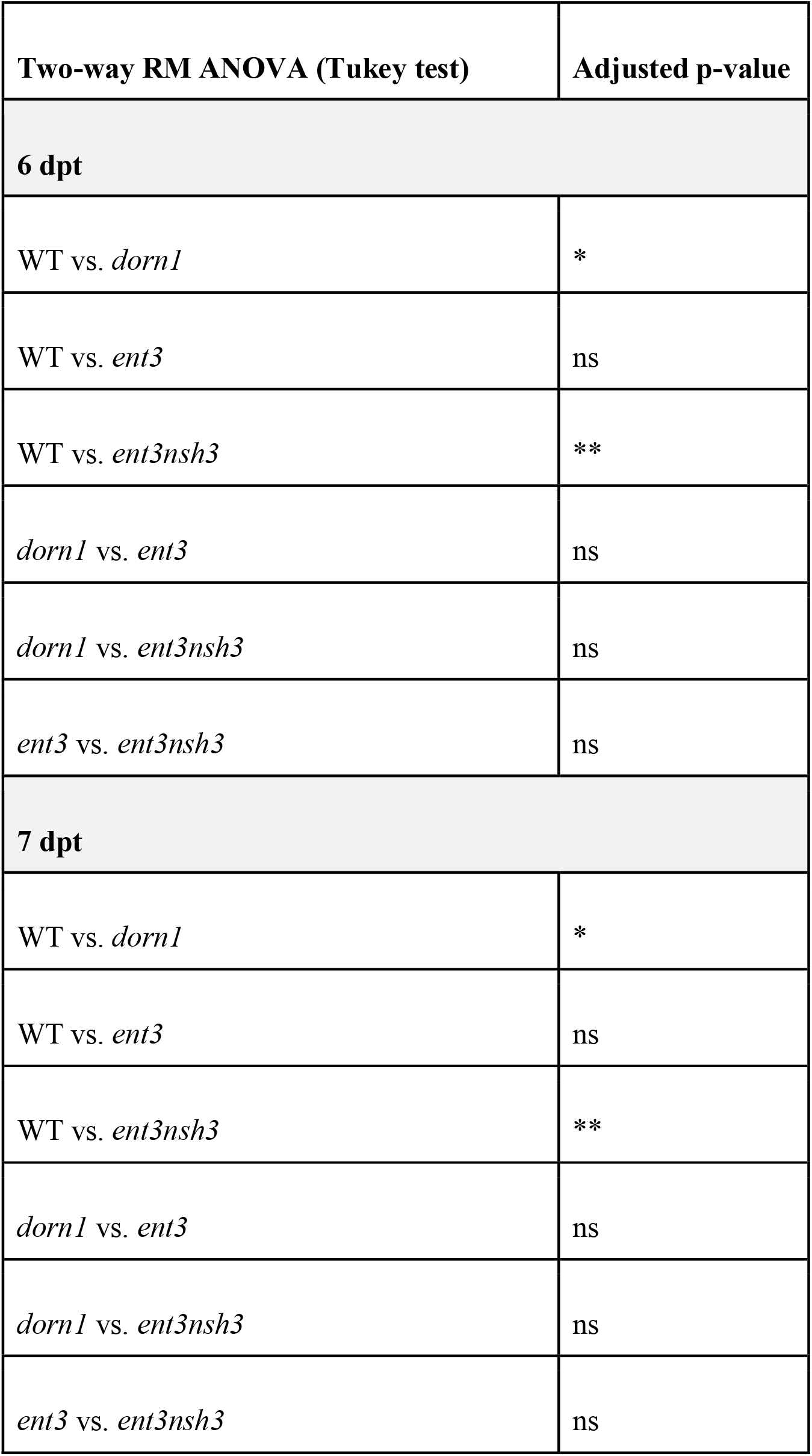
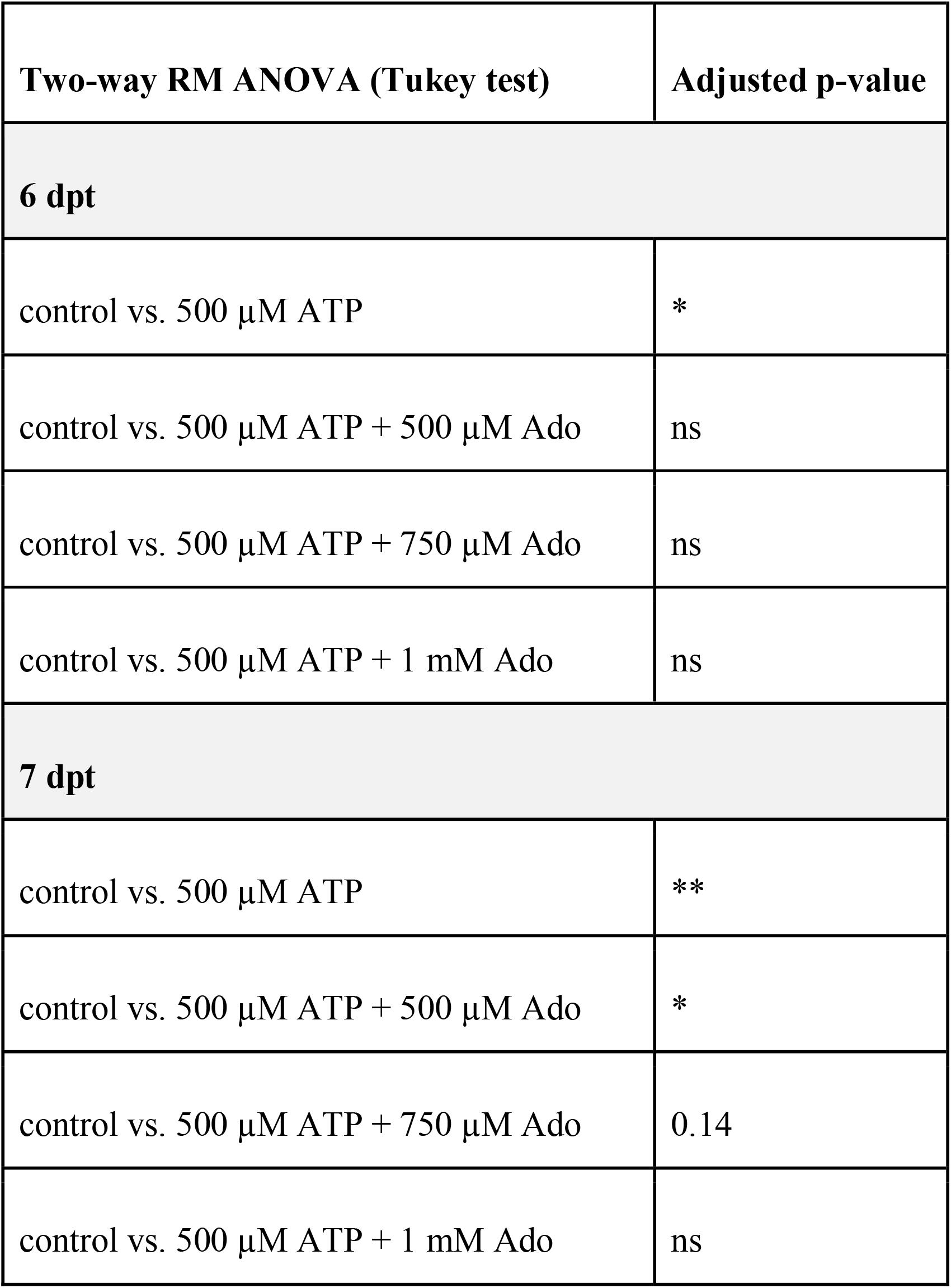
Statistical analysis of root vascular penetrations upon Fo5176 pSIX1::GFP infection. **Table S2A (linked to Figure 1D).** **Table S2B (linked to Figure 2A).** **Table S2C (linked to Figure S2B).**

